# *Citrobacter rodentium* infection activates colonic lamina propria group 2 innate lymphoid cells

**DOI:** 10.1101/2025.01.02.631085

**Authors:** Rita Berkachy, Vishwas Mishra, Priyanka Biswas, Gad Frankel

## Abstract

Group 3 innate lymphoid cells (ILC3s) play a major role in protecting against infection with the enteric mouse pathogen *Citrobacter rodentium,* used to model infections with enteropathogenic and enterohaemorrhagic *Escherichia coli*. ILC3s-secreted IL-22, shown to be indispensable for protection against *C. rodentium* infection, induces secretion of IL-18, antimicrobial peptides and nutritional immunity proteins as well as activation of tissue regeneration processes. While ILC2s have traditionally been associated with immune responses to helminth infection and allergic inflammation via the production of type 2 cytokines (e.g. IL-4, IL-5, IL-9 and IL-13), more recently they have been implicated in protection against *Clostridium difficile* and *Helicobacter pylori* infections. Here we show that colonic lamina propria ILC2s proliferate in response to *C. rodentium* infection and secrete IL-4, IL-5 and IL-13, which are involved in maintenance of the intestinal barrier function, tissue repair and mucus secretion. When stimulated with IL-18, colonic ILC2s from uninfected naïve mice secreted type 2 cytokines. Injection of IL-18 binding protein (IL18BP), at 2- and 3-days post *C. rodentium* infection, blocked activation of ILC2s. These data show that ILC2s are activated in response to infection with an enteric Gram-negative pathogen, where stimulation with IL-18 plays a role in inducing proliferation and secretion of type 2 cytokines.

**Author Summary:** While group 3 innate lymphoid cells (ILC3s) play a key role in protecting from bacterial infections, ILC2s are mainly associated with immune responses to helminth infection. Here we investigated if ILC2s are activated in responses to infection with the enteric mouse pathogen *Citrobacter rodentium.* We show that in infected mice, gut ILC2s expand and secreted type 2 cytokines. ILC2 isolated from uninfected mice were activated by IL-18. Consistently, administration of IL-18 binding protein into *C. rodentium*-infected mice inhibited ILC2 activation. These findings suggest that gut ILC2s are activated by Gram negative enteric pathogens, which is mediated in part by IL-18.

## Introduction

*Citrobacter rodentium* is the etiologic agent of transmissible murine colonic crypt hyperplasia (CCH) (1, 2). It is an enteric extracellular Gram-negative pathogen, which serves to model infections with enteropathogenic *Escherichia coli* (EPEC) and enterohemorrhagic *E. coli* (EHEC) (3, 4). To bind intestinal epithelial cells (IECs) and proliferate, *C. rodentium* employs a type III secretion system (T3SS), which injects effector proteins directly into IECs (5). Following injection, the effectors form an intracellular network which subverts signal transduction in the host cell, including trafficking and tight junctions, cell death pathways, NF-kB, JNK/p38 and caspase-4/11, -8, and -9 (6–8). Infection of *C. rodentium* (as EPEC and EHEC) is unique, as the pathogen injects its own receptor, Tir, into infected IECs (9). Binding of the outer membrane protein intimin to Tir leads to intimate bacterial attachment, effacement of the brush border microvilli, induction of localised actin polymerisation and formation of a pedestal-like structure under the attached bacteria (10).

The *C. rodentium* infection cycle is divided into four phases (11, 12): An establishment phase (1-3 days post infection (dpi)), where the pathogen colonises the caecal lymphoid patch (13). An expansion phase (4-8 dpi), initially characterised by activation of group 3 innate lymphoid cells (ILC3s)-secreting IL-22 (4 dpi), which is believed to be indispensable for survival (14–17). Binding of IL-22 to the IL-22 receptor in IECs triggers expression of IL-18, antimicrobial peptides (e.g. Reg3γ and Reg3β) and nutritional immunity (e.g. LCN-2 and calprotectin) promoting epithelial barrier resistance (17–19). This is followed (6-8 dpi) by rapid *C. rodentium* proliferation, adherence to IECs in the distal colon and development of CCH (12). A steady-state phase (9-12 dpi), characterised by high and stable *C. rodentium* shedding and a switched of IL-22 production from ILC3s to CD4 T cells, which induces sustained STAT3 activation in both superficial and crypt IECs, limiting bacterial dissemination (17, 20). A clearance phase (from 12 dpi), when *C. rodentium* is rapidly cleared via opsonisation with IgG and phagocytosis by neutrophils (20, 21).

ILC2s have been implicated in immune responses to extracellular helminth infections via the secretion of type 2 cytokines IL-4, IL-5, IL-9 and IL-13 (22, 23). However, recent studies have shown that secretion of IL-5 and IL-13 from ILC2s protects mice against toxin-mediated epithelial damage during *Clostridium difficile* infection (CDI) (24, 25). Administration of IL-13 protected mice from CDI-induced weight loss and severe disease, while anti-IL-13 exacerbated infection outcomes (26). During *Helicobacter pylori* infection, ILC2-secreting IL-5 influenced B cell responses and enhanced IgA coating of the bacteria, helping control the infections (27).

ILC2s are mainly activated by IL-25, IL-33 and TSLP which are released from epithelial and stromal cells (28). More recently, IL-18 has been shown to activate resident ILC2s in the skin (29). IL-33 has been shown to be expressed by IECs in *C. rodentium*-infected mice and higher *C. rodentium* faecal colony forming units (CFUs) were recovered from *Il33^-/-^* compared with WT control mice at 14 dpi (30). In contract, *Il25^-/-^* presented only subtle difference in infection outcomes compared to WT mice (31). The role of TSLP in *C. rodentium* infection is not known. However, IL-18 has been shown to have a protective role during *C. rodentium* infection as infection of *Il18^-/-^*resulted in greater bacterial load and exacerbated histopathology (32, 33). While IL-33 and IL-18 have been shown to play a role in *C. rodentium* infection, it is not known if they signal to colonic lamina propria ILC2s. In this study we found that the ILC2 population expanded upon *C. rodentium* infection at 4 dpi, when ILC3s have already been activated (14, 15, 34). ILC2s isolated from *C. rodentium*-infected mice produced elevated amounts of IL-5, IL-4 and IL-13. Stimulation of ILC2s, isolated from the colons of naïve mice, with IL-18 or IL-33 resulted in proliferation and secretion of IL-5, IL-9, and IL-13. These findings suggest that in parallel to ILC3s, ILC2s are also activated during the expansion phase of *C. rodentium* infection.

## Results

### Colonic ILC2s proliferate in response to *C. rodentium* infection

The aim of this study was to determine whether ILC2 respond to enteric Gram-negative bacterial infections. First, we infected C57BL/6 mice with *C. rodentium* by oral gavage and enumerated faecal CFU at 4 dpi, when it is known that ILC3s-derived IL-22 activates IECs (17). This revealed shedding at ca. 10^8^ CFU/g of stool (Fig. 1A). We then measured crypt length in H&E-stained paraffin-embedded colonic sections, which showed no significant increase in CCH compared to the uninfected control mice (Fig. 1B and 1C). In contrast, the colon length of *C. rodentium-*infected mice was significantly shorter than PBS-dosed mice and faecal myeloperoxidase (MPO), a neutrophil-derived biomarker, was secreted from infected tissue explants at significantly higher level than the control (Fig. 1D-F). Once we confirmed that the mice were infected and responded to the pathogen, we investigated colonic lamina propria ILC2s; mice dosed with PBS were used as controls. We assessed the numbers and proportions of ILC2s by flow cytometry, using a combination of lineage markers, CD45, KLRG1, CD127 and the transcription factor GATA-3 (Fig. 1G). This revealed that the number and proportion of ILC2s increased significantly in the *C. rodentium*-infected mice (Fig. 1H).

**Figure 1.**
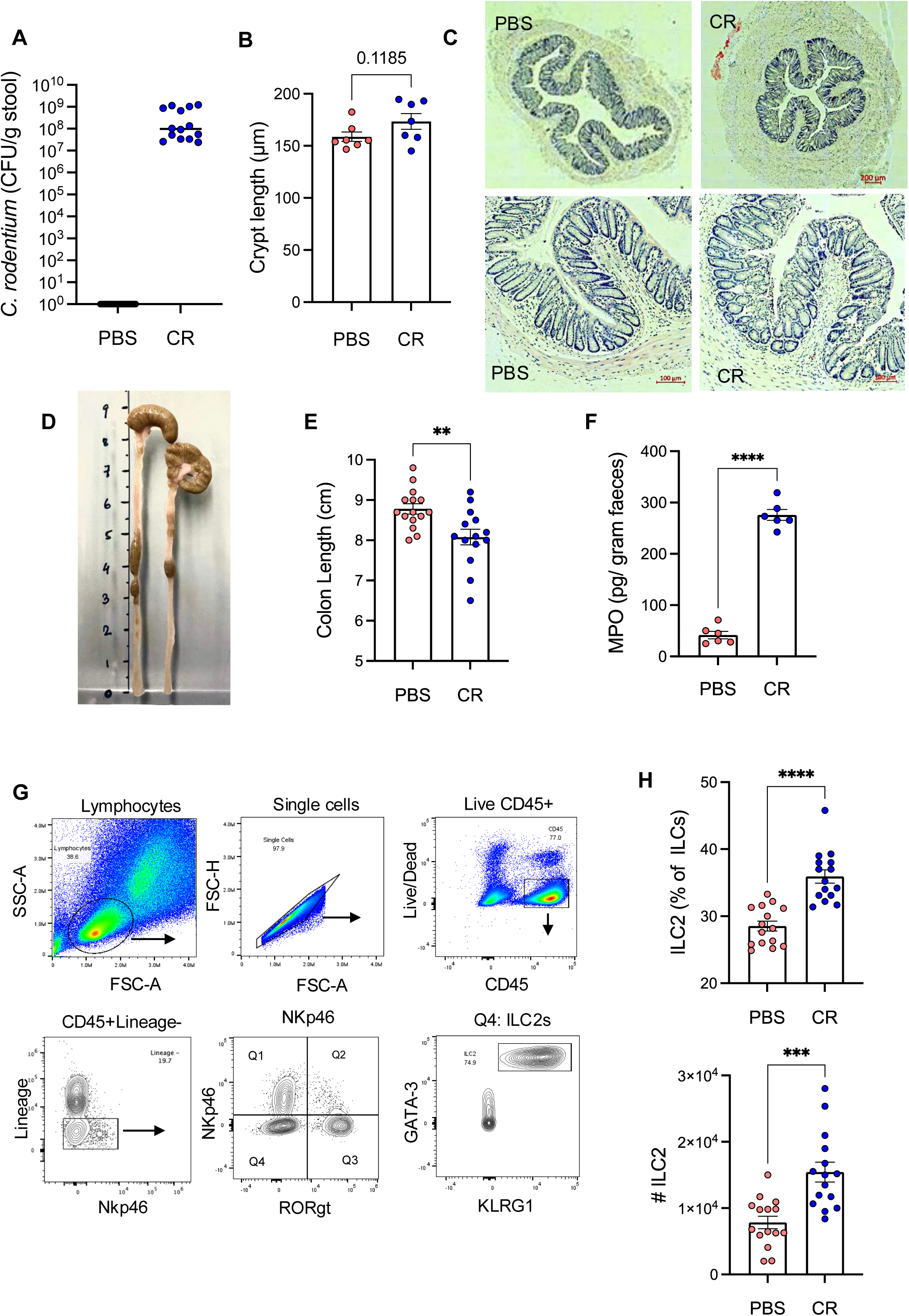
Colonic lamina propria ILC2s proliferate in response to *C. rodentium* at 4 dpi. **A.** Faecal *C. rodentium* (CR) at 4 dpi. Mock infected (PBS) were used as controls. **B.** Crypt lengths measured in H&E-stained colon sections. **C.** Representative H&E sections from the distal colons of uninfected (PBS) and infected mice at 4 dpi (scale bar 100 μm). **D.** Representative and **E.** quantitative colon length. **F.** Faecal MPO levels measured by ELISA. **G.** Flow cytometry gating strategy used to identify ILC2s from mice colonic lamina propria cells. Debris (SSC-A vs. FSC-A) and doublets (FSC-H vs. FSC-A) were excluded, and live/dead discrimination was determined using the LIVE/DEAD fixable blue dead cell stain kit (FSC-A vs. Live/Dead). Total ILCs were defined as CD45^+^NKp46^+/-^Lineage^-^ [CD3/B220/CD19/TER119/Gr-1/CD5/CD11c/CD4/Nk1.1]. Expression of the CD127, KLRG1 and GATA3 markers were used to identify ILC2 (CD127^+^KLRG1^+^GATA3^+^). **H.** Numbers and frequency of ILC2s increased upon infection. dpi: days post infection. Data shown are mean ± SEM from three independent experiments.

### *C. rodentium* infection induces type 2 cytokines secretion

To determine their cytokine profile, we sorted ILC2s from pooled colons of uninfected and *C. rodentium*-infected mice (Fig. 2A). Upon stimulation with PMA/ionomycin (without protein transport inhibitors), the levels of the type 2 cytokines were measured by ELISA. This revealed that ILC2s isolated from *C. rodentium*-infected mice secreted elevated amounts of IL-4, IL-5 and IL-13 (Fig. 2B). When protein transport inhibitor cocktail was present in the stimulation media, IL-4^+^, IL-5^+^ and IL-13^+^ ILC2 populations were higher in *C. rodentium* infected samples (Fig. 2 C-D), suggesting that colonic lamina propria ILC2s not only proliferated but were also activated in *C. rodentium*-infected mice.

**Figure 2.**
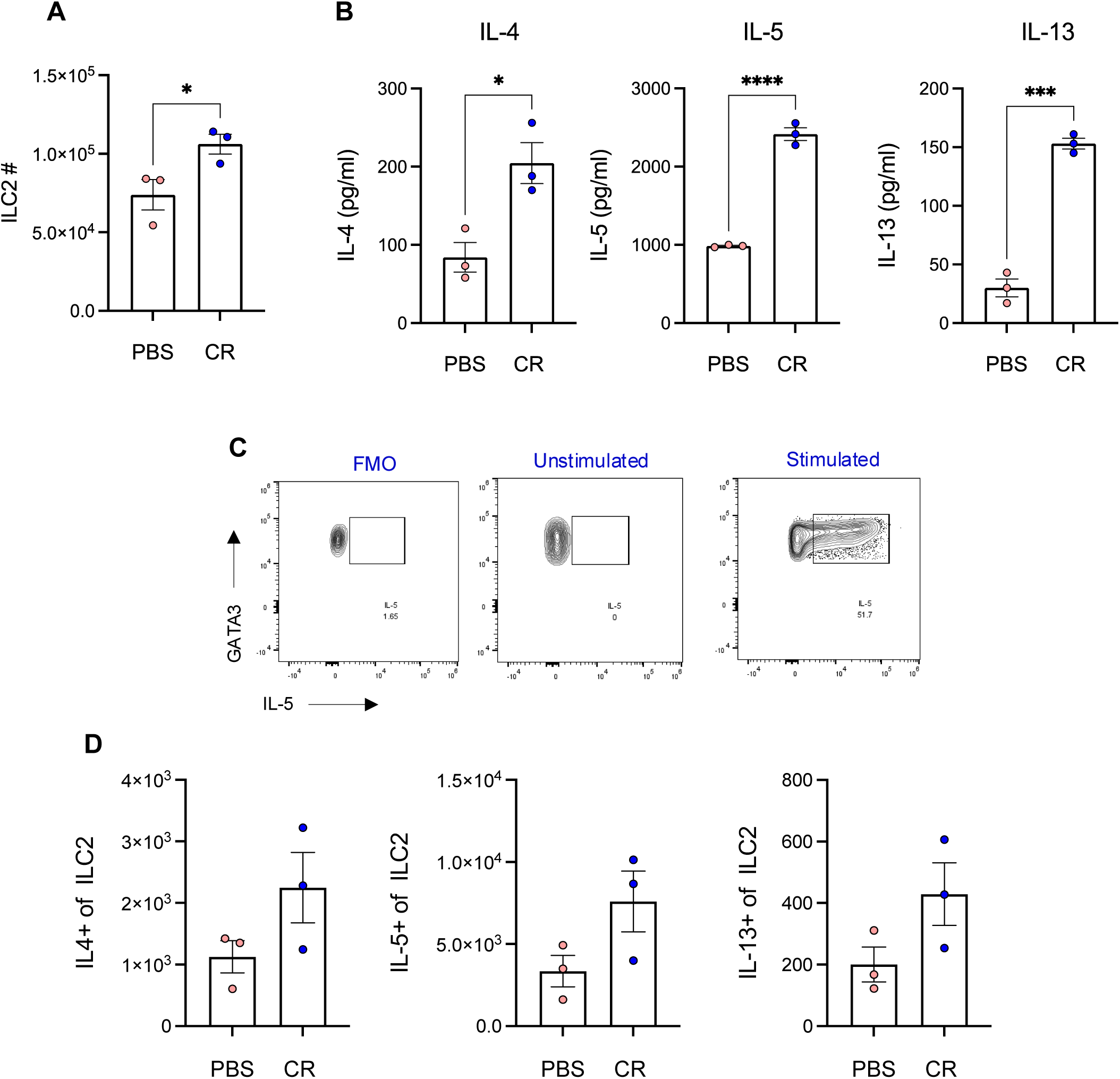
ILC2 cytokine profile during *C. rodentium* infection. **A.** Total ILC2 numbers sorted from pooled colons of uninfected and infected mice. **B.** IL-4, IL-5 and IL-13 levels in the supernatant from the same sorted cells measured by ELISA. **C.** Representative flow cytometry plots of cytokine expression from fluorescent-minus-one (FMO), unstimulated and stimulated samples. **C.** Total number of IL-4^+^, IL-5^+^ and IL-13^+^ of sorted ILC2 cells from colons pooled from uninfected and infected mice. Cells were stimulated for 4 h in the presence of protein transport inhibitor cocktail to assess cytokine intracellularly by flow cytometry. Data shown are mean ± SEM from three independent experiments.

### Colonic ILC2s proliferate in response to IL-18

We next evaluated if any of the ILC2-activating cytokines are expressed in IECs 4 days post *C. rodentium* infection. We extracted mRNA from whole colonic tissue and performed RT-qPCR. This revealed that while *Il25* expression did not increase and *TLSP* was not detected, expression of *Il18 and Il33* was elevated in *C. rodentium*-infected mice (Fig. 3A). Moreover, ELISA revealed that IL-18 levels, secreted from tissue explants, were significantly higher in the infected samples (Fig. 3B).

**Figure 3.**
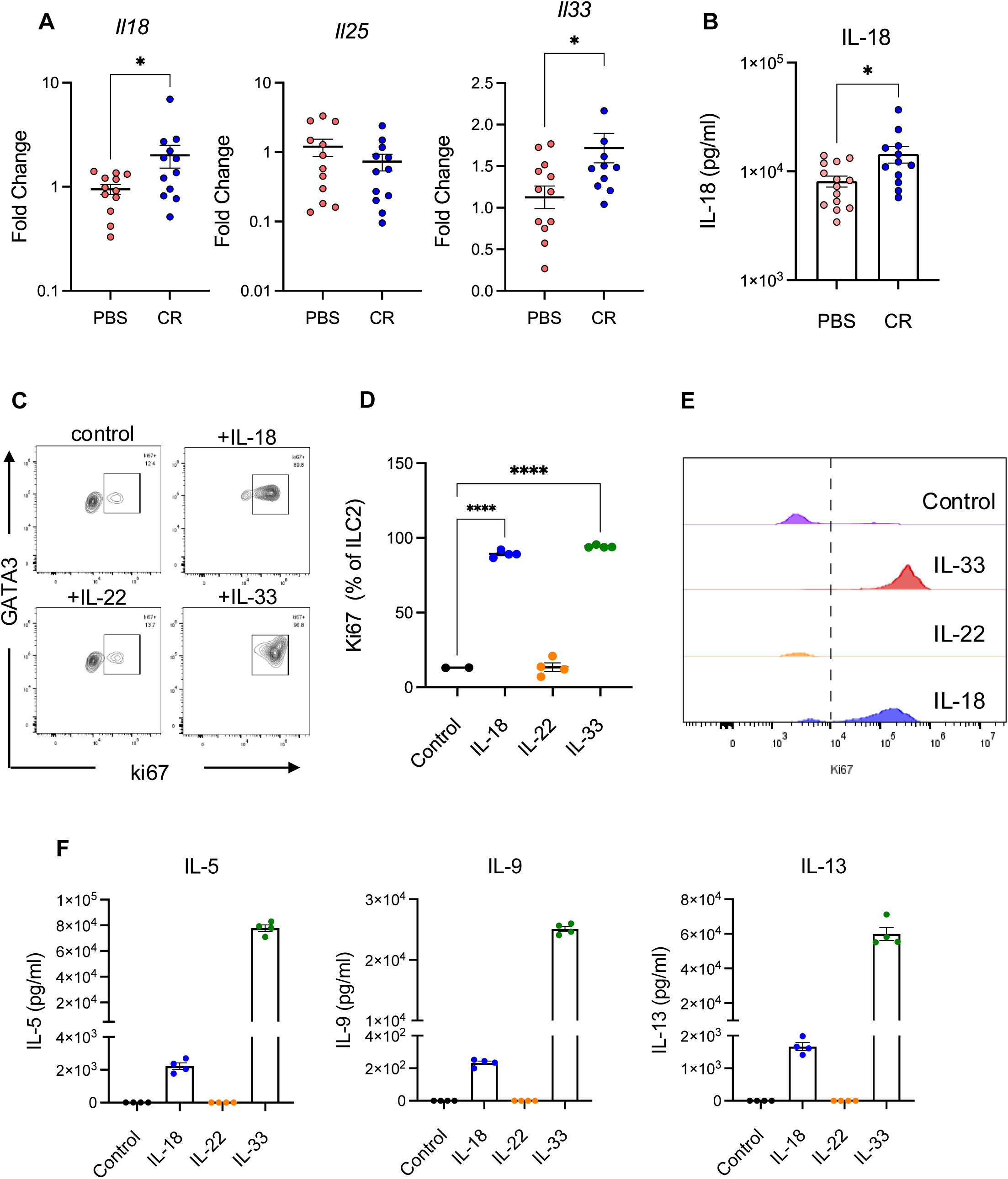
Colonic lamina propria ILC2s proliferate in response to IL-18. **A.** RT-qPCR analysis of *Il18, Il25* and *Il33* transcript expression in enriched IECs from colons of CR-infected and PBS-dosed mice at 4 dpi. Each data point represents one mouse. **B.** IL-18 levels secreted from explant colonic tissue and detected by ELISA. **C.** Representative flow cytometry plots and **D.** Summary data for Ki67 expression (represented as percentage of ILC2) by FACS sorted colonic lamina propria ILC2s. Cells were cultured for 3 days *in vitro* with recombinant IL-2 and IL-7 (control) in addition to IL-18, IL-22 or IL-33. **E.** Histogram showing positive and negative peaks of proliferative ILC2s in response to different cytokines. **F.** IL-5, IL-9 and IL-13 levels in the supernatant from cultured ILC2s measured by ELISA. Cells pooled from n = 10 mice per experiment. Data shown are mean ± SEM from four independent experiments.

We investigated if, similarly to skin ILC2s, IL-18 could also stimulate colonic lamina propria ILC2s (29). To this end, ILC2s sorted from the colons of naïve mice were cultured for 3 days in complete RPMI medium containing recombinant murine IL-2 (2 ng/ml) and IL-7 (5 ng/ml), in addition to IL-18 (100 ng/ml). IL-33 and IL-22 (100 ng/ml) were included as positive and negative controls, respectively. Quantification of the proliferation marker Ki67 revealed that ILC2s proliferated in response to IL-33 and IL-18, but not IL-22 (Fig. 3C-E). Analysing supernatants of stimulated ILC2s for cytokine production by ELISA showed that only IL-18 and IL-33 triggered secretion of IL-5, IL-9, and IL-13 (Fig. 3F), while IL-4 was not detected.

Finaly, to test if IL-18 is involved in activation of ILC2 *in vivo*, *C. rodentium*-infected mice were treated with IL-18 binding protein (IL-18BP) intraperitoneally at 2 and 3 dpi (Fig. 4A). Treated mice presented a significantly lower number and frequency of ILC2s compared to the vehicle control group (Fig. 4B). These findings suggest that IL-18 stimulates colonic lamina propria ILC2s during *C. rodentium* infection.

**Figure 4.**
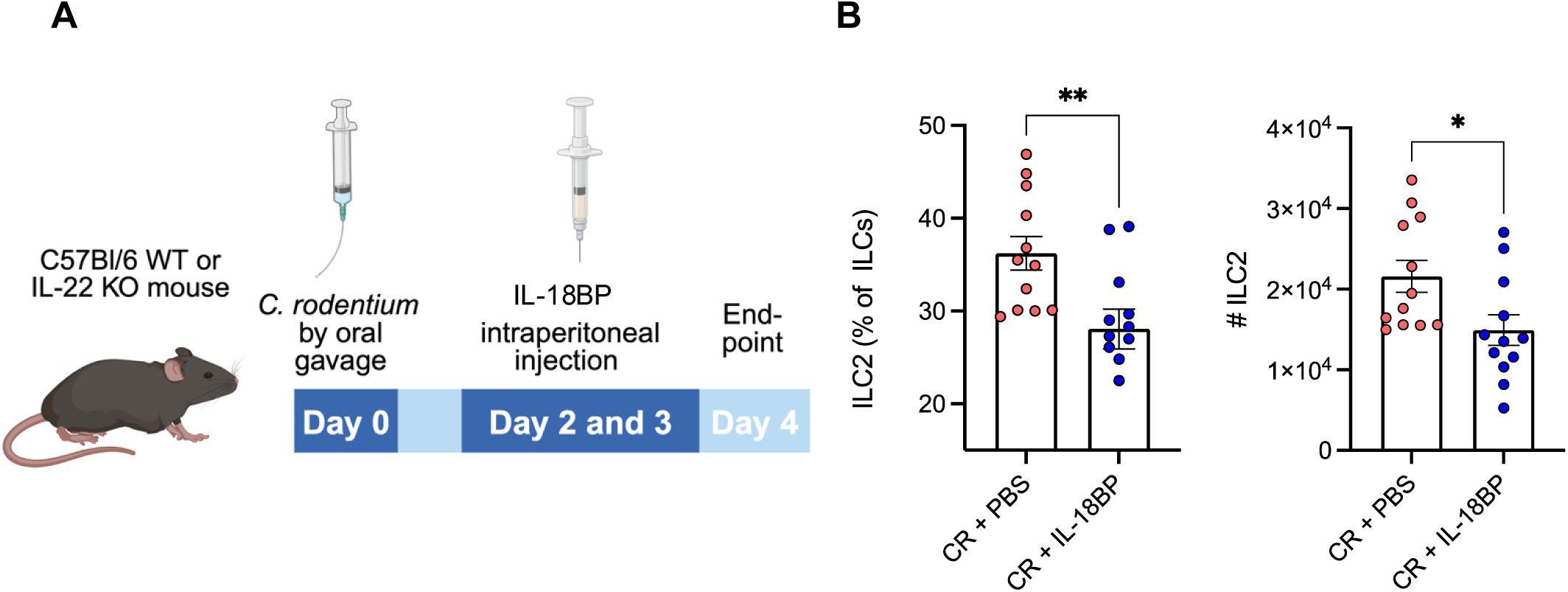
IL-18 restored the stimulation of ILC2s in IL-22 KO mice. **A.** Experimental plan. **B.** Frequency and absolute number of ILC2s is reduced in the colon of CR infected C57BL/6J mice when treated with IL-18BP. Data shown are mean ± SEM from three independent experiments.

## Discussion

ILC3s are one of the first responders to gut perturbation and infection with extracellular enteric pathogens, while ILC2s are mainly associated with helminth infection (22, 23). More recently, ILC2s have been implicated in colonic CDI and gastric *H. pylori* infection (24, 25, 27). In this study we provide first evidence that colonic lamina propria ILC2s proliferate and produce type 2 cytokines in response to *C. rodentium* infection. While we do not yet know the role ILC2s, and their secreted cytokines, play during the infection, our data suggest that ILC2s are activated in the colonic lamina propria in response to Gram-negative bacterial pathogens.

ILC3s are known to be activated at 4 dpi, when *C. rodentium* sparsely and sporadically colonises the apex of the colonic crypts (15). Here we show that at 4 dpi, the colonic tissue shows morphological signs of inflammation, i.e. shortening of the colon and secretion of MPO and IL-18. This is accompanied by a significant increase in population of colonic lamina propria ILC2s. Of note, of the major stimulators of ILC2s, we did not detect induction of *Il25* expression, while *TLSP* was undetected in the large intestine. In contrast, expression of *Il33* and *Il18* was induced in *C. rodentium*-infected mice. In terms of type 2 cytokines, stimulation of ILC2s isolated from infected mice, enhanced the production of IL-4, IL-5 and IL-13. Conversely, upon stimulation with IL-33 and IL-18 of ILC2s isolated the colons of naïve mice and maintained in vitro for 3 days, we detected secretion of IL-5, IL-9, and IL-13; IL-4 was not detected. This shows important differences in ILC2 responses to IL-18 and IL-33 stimulation dependent on whether they were isolated from infected or naïve mice.

While their role is not yet known, it is possible that IL-4, IL-5 and IL-13 participate in mucus secretion and epithelial repair in early responses to *C. rodentium* colonisation in order to maintain barrier function. Mice deficient in IL-13 exhibit impaired barrier function and increased susceptibility to intestinal pathogens (35–37). During helminth infections, IL-13 produced by ILC2s drives goblet cell proliferation, mucus secretion and smooth muscle contractility, contributing to parasite expulsion (38). On the other hand, IL-5 is associated with the recruitment of eosinophils but can also enhance IgA production controlling bacterial spread in *H. pylori* infected mice (27, 39, 40). IL-4, however, plays a role in regulating IgE responses in B cells and the differentiation of alternatively activated M2 macrophages (41). In addition, IL-4, secreted from CD4+ T cells, has been shown to increase mucus production in *C. rodentium*-infected mice (42). The significance of ILC2 activation during *C. rodentium* infection could lie in production of IL-4 and IL-13 to drive mucus production, in addition to IL-5 which can act synergistically with IL-4 to influence or enhance an antibody response, promoting bacterial clearance (43). Moreover, ILC2s have been shown to synthesise and release acetylcholine (ACh) during parasitic nematode infection (44). Depleting ACh production in T cells led to a higher bacterial burden and increased levels of IL-1β, IL-6 and TNFα in the gut of *C. rodentium* infected mice (45). Therefore, ILC2s may play a supportive role by limiting tissue damage and promoting epithelial repair rather than directly controlling bacterial growth through type 2 cytokine secretion. This dual role of ILC2s in inflammation and repair is supported by recent work highlighting their plasticity and responsiveness to diverse cytokine milieus (46).

We show that IL-18 and IL-33, detected at 4 dpi, stimulate colonic ILC2s proliferation and cytokine production. In contrast, treating mice with IL-18BP at days 2 and 3 post-*C. rodentium* infection inhibited ILC2s expansion, providing direct evidence that IL-18 plays a key role in ILC2s activation during infection. While IL-33 is a well-established ILC2s activator (47, 48), our findings add to the emerging evidence that IL-18, traditionally associated with type 1 responses, can activate ILC2s under certain conditions. Skin ILC2 subsets have been shown to be activated dominantly by IL-18 via the IL-18R1 receptor, which is highly expressed in skin ILC2s and can be detected on lung and bone marrow, but not on small intestine ILC2s (29). Our data suggest that the IL-18R1 receptor is also expressed on ILC2s in the large intestine.

Taken together, this study shows that colonic ILC2s proliferate and secrete type 2 cytokines in response to *C. rodentium* infection, driven, at least in part, by IL-18 and IL-33. Further studies using genetically modified mice and neutralisation of type 2 cytokines are now needed to determine the functional role of ILC2s in host responses to *C. rodentium* and how widespread this response is in other bacterial enteric infections.

## Materials and methods

### Mice and ethics statement

Female C57Bl/6J mice of 6-10 weeks age were purchased from Charles River UK. All mice were maintained at the Central Biomedical Services (CBS) facility at Imperial in pathogen-free conditions at 20-22 °C and 30-40 % humidity on 12 h light/dark cycle. All animal procedures were conducted in accordance with Animals Scientific Procedures Act 1986 and U.K. Home Office guidelines and reviewed by the Imperial College Animal Welfare Ethical Review Body (AWERB). The in vivo procedures are covered by a home office licence, number PP7392693.

### C. rodentium infection

*C. rodentium* ICC169 cultures were prepared in 15 ml LB supplemented with 50 μg/μL nalidixic acid and grown at 37 °C overnight. The cultures were centrifuged, resuspended with 1.5 ml phosphate-buffered saline (PBS) and each mouse received 200 μl by oral gavage (12). Uninfected mice (mock infected) received 200 μl of PBS. To quantify the colony forming unit (CFU) per gram of stool, stool samples were homogenised, diluted and plated onto agar plates and viable bacteria were counted as previously described (12). Infected mice that did not reach the threshold of 1 x 10^7^ CFU/g of stool were excluded from further experimental processing and analysis.

### Administration of IL-18 binding protein

Recombinant human IL-18 binding protein (rhIL-18BP, Cat No. 119-BP, R&D Systems) was freshly prepared in PBS and given to mice by intraperitoneal injection, in a volume of 0.1 ml on days 2 and 3 p.i. at a dose of 5 mg/kg and 1 mg/kg respectively. Control groups received PBS only.

### Cytokines and antibodies

Recombinant murine cytokines were purchased from PeproTech (IL-2, IL-7, IL-22, and IL-33) or R&D Systems (IL-18 and TSLP). All cytokines were lyophilised in PBS and stored at -80 °C. The following antibodies were purchased from Biolegend or BD Biosciences and used for flow cytometry: Fc Block (anti-mouse CD16/CD32 mAb, clone 2.4G2, BD Biosciences), CD3 (145-2C11), B220 (RA3-6B2), CD19 (6D5), TER119 (TER-119), Gr-1 (RB6-8C5), CD5 (53-7.3), FcεRI (Marl), CD11c (N418), F4/80 (BM8), CD45 (30F11), CD90.2 (30-H12), Nkp46 (29A1.4), CD4 (RM4-5), CD127 (A7R34), KLRG1 (2F1), NK1.1 (PK136), GATA3 (L50-823), RORγt (Q31-378), IL-4 (11B11), IL-5 (TRFR5), IL-13 (ebio13A) and Ki67 (SolA15).

### Isolation of lamina propria immune cells and IECs from mouse colons

Lamina propria cells were isolated from mouse colons as previously described (49). Briefly, cleaned colons were cut longitudinally then incubated for 20 min at 37 °C shaking in Hanks’ balanced salt solution (HBSS; Ca^2+^ and Mg^2+^ free) supplemented with 2% FBS, 10 mM EDTA and 1 mM DTT. Residual tissues were digested at 37°C for ∼40-50 min in RPMI 1640 containing 62.5 μg/mL Liberase, 50 μg/mL DNase I and 2 % FBS. Mononuclear cells were isolated with 40-80% Percoll gradient and washed twice.

### Flow cytometry and cell sorting

For the analysis of cytokine expression, isolated cells were stimulated with 1x cell stimulation cocktail (Cat No. 00-4970, eBioscience) for 4 h at 37°C / 5% CO_2_ in the presence or absence of 1x protein transport inhibitor cocktail (Cat No. 00-4980, eBioscience).

To stain for extracellular markers, single-cell suspensions or stimulated cells were plated in a 96 well V bottomed plate and stained for 10 min with LIVE/DEAD fixable blue (Thermo Fisher, diluted in D-PBS) to detect and exclude dead cells from subsequent analysis. Cells were then treated for 20 min with Fcγ receptor block followed by surface marker staining using fluorophore-conjugated monoclonal antibodies (BioLegend, BD Biosciences or eBioscience). All incubations were performed at 4°C in the dark, unless otherwise stated. Negative (unstained and live/dead) along with fluorescent-minus-one (FMO) controls were considered to estimate background fluorescence.

For intracellular staining, cells were first fixed for 20-30 min at room temperature in the dark, using the eBioscience Forkhead box protein 3 (Foxp3)/transcription factor fixation buffer set to stain for transcription factors or the IC fixation buffer for the detection of intracellular cytokines. Fixed cells were then stained for 1 h with intracellular antibody cocktail diluted in permeabilisation buffer. Cells were then washed prior to analysis on an Aurora flow cytometer (Cytek Biosciences).

For ILC2 sorting, lineage negative population (CD3ε, CD4, CD8α, CD5, CD19, CD11c, Gr1, F4/80, FcεRIa, NK1.1, CD11b, and TER119) were excluded by Magnetic Cell sorting system (MACS, Miltenyi Biotec) from single-cell suspension. MACS separated cells were blocked with anti-CD16/CD32 for 20 min on ice then stained with CD45, CD127, KLRG1, CD90.2 and lineage cocktail antibodies for 1 hour on ice. After staining, cells were washed and resuspended in DAPI 5 min prior to sorting and ILC2 cells were sorted on a FACSAria III cell sorter (BD Biosciences).

### ILC2 in vitro culture and proliferation assay

For each experimental repeat, colons from 15 mice were pooled and sorted as a single sample, and isolated ILC2 cells were split equally between different conditions. ILC2s were cultured for 3 days at 37°C / 5% CO_2_ in U-bottom 96-well plate in complete RPMI medium containing recombinant murine IL-2 (2 ng/ml), IL-7 (5 ng/ml) and either IL-18, IL-22 or IL-33 (100 ng/ml). Cells were re-stained using the extracellular ILC2 marker panel followed by intranuclear Ki67 staining, and cell proliferation was assessed by flow cytometry.

### RNA isolation, reverse transcription and quantitative real-time PCR

Total RNA was extracted from whole tissue using the RNeasy Mini Kit (QIAGEN) and converted to cDNA using iScript cDNA synthesis kit (BIO-RAD) in accordance with the manufacturer’s instructions. The RT-qPCR reactions were carried out using the PowerUP SYBR green PCR master mix (Applied Biosystems) and run in duplicates in a 7500 Fast RT-PCR thermocycler (Applied Biosystems). Relative gene expression of *Il18*, *Il25*, *Il33* and *Tslp* was calculated by the comparative cycle threshold (Ct) method 2^-ΔΔCt^, using *Hprt, Actb* and *Gapdh* as housekeeping genes. Primer sequences were as follows: *Il18* forward: GACTCTTGCGTCAACTTCAAGG; *Il18* reverse: CAGGCTGTCTTTTGTCAACGA; *Il25* forward: : ACAGGGACTTGAATCGGGTC; *Il25* reverse: TGGTAAAGTGGGACGGAGTTG; *Il33* forward: TCCAACTCCAAGATTTCCCCG; *Il33* reverse: CATGCAGTAGACATGGCAGAA; *Tslp* forward: : ACGGATGGGGCTAACTTACAA and *Tslp* reverse: AGTCCTCGATTTGCTCGAACT.

### Cytokine analysis by ELISA

The levels of IL-4, IL-5, IL-9, IL-13 and IL-18 were determined using Duoset ELISA kits (R&D Systems). For tissue explant, 1 cm colon fragments were transferred to a 96-well plate containing RPMI buffer supplemented with 10 % faetal bovine serum (FBS), 100 U/ml penicillin, 100 µg/ml streptomycin and incubated in a 37°C humidified 5 % CO_2_ incubator for 24 h. Supernatants from cultured ILC2 or colonic explants were collected, and cytokines were analysed by ELISA (R&D Systems) according to the manufacturer’s instructions.

### Statistical analysis

Flow cytometry data were analysed using FLowJo software (Tree Star). Graphs and statistical tests were carried out using GraphPad Prism 10.2.3 software and data were expressed as mean ± SEM. Statistical differences were calculated using the unpaired Mann-Whitney Student’s t test for two groups and one-way ANOVA for 3 or more groups. A p value of <0.05 is considered significant, *p<0.05, **p<0.01, ***p<0.001.

Custom images of mice in Fig. 4C and graphical abstract were made with Biorender (created with BioRender.com).

## Acknowledgments

We thank Jessica Rowley and Larissa Zarate Garcia for their technical assistance with flow cytometry and cell sorting. This study was supported by grants from The Wellcome Trust (224282/Z/21/Z) and the Medical Research Council (MRC) (MR/R02671).

## References

1. Barthold SW, Coleman GL, Bhatt PN, Osbaldiston GW, Jonas AM. The etiology of transmissible murine colonic hyperplasia. Lab Anim Sci. 1976;26(6 Pt 1):889–94.

2. Luperchio SA, Newman JV, Dangler CA, Schrenzel MD, Brenner DJ, Steigerwalt AG, et al. *Citrobacter rodentium*, the causative agent of transmissible murine colonic hyperplasia, exhibits clonality: synonymy of *C. rodentium* and mouse-pathogenic *Escherichia coli*. J Clin Microbiol. 2000;38(12):4343–50.

3. Mundy R, MacDonald TT, Dougan G, Frankel G, Wiles S. *Citrobacter rodentium* of mice and man. Cell Microbiol. 2005;7(12):1697–706.

4. Mallick EM, McBee ME, Vanguri VK, Melton-Celsa AR, Schlieper K, Karalius BJ, et al. A novel murine infection model for Shiga toxin-producing *Escherichia coli*. The Journal of clinical investigation. 2012;122(11):4012–24.

5. Deng W, Marshall NC, Rowland JL, McCoy JM, Worrall LJ, Santos AS, et al. Assembly, structure, function and regulation of type III secretion systems. Nat Rev Microbiol. 2017;15(6):323–37.

6. Wong AR, Pearson JS, Bright MD, Munera D, Robinson KS, Lee SF, et al. Enteropathogenic and enterohaemorrhagic *Escherichia coli*: even more subversive elements. Mol Microbiol. 2011;80(6):1420–38.

7. Pearson JS, Hartland EL. The inflammatory response during enterohemorrhagic *Escherichia coli* infection. Microbiol Spectr. 2014;2(4):EHEC-0012-2013.

8. Ruano-Gallego D, Sanchez-Garrido J, Kozik Z, Nunez-Berrueco E, Cepeda-Molero M, Mullineaux-Sanders C, et al. Type III secretion system effectors form robust and flexible intracellular virulence networks. Science. 2021;371(6534).

9. Kenny B, DeVinney R, Stein M, Reinscheid DJ, Frey EA, Finlay BB. Enteropathogenic *E. coli* (EPEC) transfers its receptor for intimate adherence into mammalian cells. Cell. 1997;91(4):511–20.

10. Frankel G, Phillips AD, Trabulsi LR, Knutton S, Dougan G, Matthews S. Intimin and the host cell--is it bound to end in Tir(s)? Trends Microbiol. 2001;9(5):214–8.

11. Mullineaux-Sanders C, Sanchez-Garrido J, Hopkins EGD, Shenoy AR, Barry R, Frankel G. *Citrobacter rodentium*-host-microbiota interactions: immunity, bioenergetics and metabolism. Nat Rev Microbiol. 2019;17(11):701–15.

12. Crepin VF, Collins JW, Habibzay M, Frankel G. *Citrobacter rodentium* mouse model of bacterial infection. Nat Protoc. 2016;11(10):1851–76.

13. Wiles S, Clare S, Harker J, Huett A, Young D, Dougan G, et al. Organ specificity, colonization and clearance dynamics in vivo following oral challenges with the murine pathogen *Citrobacter rodentium*. Cell Microbiol. 2004;6(10):963–72.

14. Serafini N, Jarade A, Surace L, Goncalves P, Sismeiro O, Varet H, et al. Trained ILC3 responses promote intestinal defense. Science. 2022;375(6583):859–63.

15. Jarade A, Di Santo JP, Serafini N. Group 3 innate lymphoid cells mediate host defense against attaching and effacing pathogens. Curr Opin Microbiol. 2021;63:83–91.

16. Zindl CL, Wilson CG, Chadha AS, Duck LW, Cai B, Harbour SN, et al. Distal colonocytes targeted by *C. rodentium* recruit T-cell help for barrier defence. Nature. 2024;629(8012):669–78.

17. Zindl CL, Witte SJ, Laufer VA, Gao M, Yue Z, Janowski KM, et al. A nonredundant role for T cell-derived interleukin 22 in antibacterial defense of colonic crypts. Immunity. 2022;55(3):494–511 e11.

18. Melchior K, Gerner RR, Hossain S, Nuccio SP, Moreira CG, Raffatellu M. IL-22-dependent responses and their role during *Citrobacter rodentium* infection. Infection and immunity. 2024;92(5):e0009924.

19. Hopkins EGD, Roumeliotis TI, Mullineaux-Sanders C, Choudhary JS, Frankel G. Intestinal epithelial cells and the microbiome undergo swift reprogramming at the inception of colonic *Citrobacter rodentium* infection. mBio. 2019;10(2).

20. Mullineaux-Sanders C, Kozik Z, Sanchez-Garrido J, Hopkins EGD, Choudhary JS, Frankel G. *Citrobacter rodentium* infection induces persistent molecular changes and interferon gamma-dependent major histocompatibility complex class II expression in the colonic epithelium. mBio. 2021;13(1):e0323321.

21. Eckmann L, Stappenbeck TS. IgG “detoxes” the intestinal mucosa. Cell Host Microbe. 2015;17(5):538–9.

22. Spits H, Mjosberg J. Heterogeneity of type 2 innate lymphoid cells. Nature reviews Immunology. 2022;22(11):701–12.

23. Kasal DN, Liang Z, Hollinger MK, O’Leary CY, Lisicka W, Sperling AI, et al. A Gata3 enhancer necessary for ILC2 development and function. Proc Natl Acad Sci U S A. 2021;118(32).

24. Frisbee AL, Saleh MM, Young MK, Leslie JL, Simpson ME, Abhyankar MM, et al. IL-33 drives group 2 innate lymphoid cell-mediated protection during *Clostridium difficile* infection. Nature communications. 2019;10(1):2712.

25. Uddin MJ, Thompson B, Leslie JL, Fishman C, Sol-Church K, Kumar P, et al. Investigating the impact of antibiotic-induced dysbiosis on protection from *Clostridium difficile* colitis by mouse colonic innate lymphoid cells. mBio. 2024;15(3):e0333823.

26. Donlan AN, Leslie JL, Simpson ME, Petri WA, Allen JE, Petri WA. IL-13 protects from *Clostridioides difficile* colitis. Anaerobe. 2024;88:102860.

27. Satoh-Takayama N, Kato T, Motomura Y, Kageyama T, Taguchi-Atarashi N, Kinoshita-Daitoku R, et al. Bacteria-induced group 2 innate lymphoid cells in the stomach provide immune protection through induction of IgA. Immunity. 2020;52(4):635–49 e4.

28. Stanbery AG, Shuchi S, Jakob von M, Tait Wojno ED, Ziegler SF. TSLP, IL-33, and IL-25: Not just for allergy and helminth infection. J Allergy Clin Immunol. 2022;150(6):1302–13.

29. Ricardo-Gonzalez RR, Van Dyken SJ, Schneider C, Lee J, Nussbaum JC, Liang HE, et al. Tissue signals imprint ILC2 identity with anticipatory function. Nat Immunol. 2018;19(10):1093–9.

30. Xiao Y, Huang X, Zhao Y, Chen F, Sun M, Yang W, et al. Interleukin-33 promotes REG3gamma expression in intestinal epithelial cells and regulates gut microbiota. Cell Mol Gastroenterol Hepatol. 2019;8(1):21–36.

31. Reynolds JM, Lee YH, Shi Y, Wang X, Angkasekwinai P, Nallaparaju KC, et al. Interleukin-17B antagonizes interleukin-25-mediated mucosaliInflammation. Immunity. 2015;42(4):692–703.

32. Liu Z, Zaki MH, Vogel P, Gurung P, Finlay BB, Deng W, et al. Role of inflammasomes in host defense against *Citrobacter rodentium* infection. The Journal of biological chemistry. 2012;287(20):16955–64.

33. Munoz M, Eidenschenk C, Ota N, Wong K, Lohmann U, Kuhl AA, et al. Interleukin-22 induces interleukin-18 expression from epithelial cells during intestinal infection. Immunity. 2015;42(2):321–31.

34. Zheng Y, Valdez PA, Danilenko DM, Hu Y, Sa SM, Gong Q, et al. Interleukin-22 mediates early host defense against attaching and effacing bacterial pathogens. Nat Med. 2008;14(3):282–9.

35. Finkelman FD, Shea-Donohue T, Morris SC, Gildea L, Strait R, Madden KB, et al. Interleukin-4- and interleukin-13-mediated host protection against intestinal nematode parasites. Immunological reviews. 2004;201:139–55.

36. Madden KB, Whitman L, Sullivan C, Gause WC, Urban JF, Jr., Katona IM, et al. Role of STAT6 and mast cells in IL-4- and IL-13-induced alterations in murine intestinal epithelial cell function. Journal of immunology (Baltimore, Md : 1950). 2002;169(8):4417-22.

37. Shea-Donohue T, Fasano A, Smith A, Zhao A. Enteric pathogens and gut function: Role of cytokines and STATs. Gut Microbes. 2010;1(5):316–24.

38. Price AE, Liang HE, Sullivan BM, Reinhardt RL, Eisley CJ, Erle DJ, et al. Systemically dispersed innate IL-13-expressing cells in type 2 immunity. Proc Natl Acad Sci U S A. 2010;107(25):11489–94.

39. Moro K, Yamada T, Tanabe M, Takeuchi T, Ikawa T, Kawamoto H, et al. Innate production of T(H)2 cytokines by adipose tissue-associated c-Kit(+)Sca-1(+) lymphoid cells. Nature. 2010;463(7280):540–4.

40. Yasuda K, Muto T, Kawagoe T, Matsumoto M, Sasaki Y, Matsushita K, et al. Contribution of IL-33-activated type II innate lymphoid cells to pulmonary eosinophilia in intestinal nematode-infected mice. Proc Natl Acad Sci U S A. 2012;109(9):3451–6.

41. Varela F, Symowski C, Pollock J, Wirtz S, Voehringer D. IL-4/IL-13-producing ILC2s are required for timely control of intestinal helminth infection in mice. Eur J Immunol. 2022;52(12):1925–33.

42. Sharba S, Navabi N, Padra M, Persson JA, Quintana-Hayashi MP, Gustafsson JK, et al. Interleukin 4 induces rapid mucin transport, increases mucus thickness and quality and decreases colitis and *Citrobacter rodentium* in contact with epithelial cells. Virulence. 2019;10(1):97–117.

43. Gustafsson JK, Navabi N, Rodriguez-Pineiro AM, Alomran AH, Premaratne P, Fernandez HR, et al. Dynamic changes in mucus thickness and ion secretion during *Citrobacter rodentium* infection and clearance. PloS one. 2013;8(12):e84430.

44. Roberts LB, Schnoeller C, Berkachy R, Darby M, Pillaye J, Oudhoff MJ, et al. Acetylcholine production by group 2 innate lymphoid cells promotes mucosal immunity to helminths. Sci Immunol. 2021;6(57).

45. Ramirez VT, Godinez DR, Brust-Mascher I, Nonnecke EB, Castillo PA, Gardner MB, et al. T-cell derived acetylcholine aids host defenses during enteric bacterial infection with *Citrobacter rodentium*. PLoS pathogens. 2019;15(4):e1007719.

46. Kotas ME, Locksley RM. Why Innate Lymphoid Cells? Immunity. 2018;48(6):1081–90.

47. Kim HY, Chang YJ, Subramanian S, Lee HH, Albacker LA, Matangkasombut P, et al. Innate lymphoid cells responding to IL-33 mediate airway hyperreactivity independently of adaptive immunity. J Allergy Clin Immunol. 2012;129(1):216–27 e1-6.

48. Klein Wolterink RG, Kleinjan A, van Nimwegen M, Bergen I, de Bruijn M, Levani Y, et al. Pulmonary innate lymphoid cells are major producers of IL-5 and IL-13 in murine models of allergic asthma. Eur J Immunol. 2012;42(5):1106–16.

49. Kim E, Tran M, Sun Y, Huh JR. Isolation and analyses of lamina propria lymphocytes from mouse intestines. STAR Protoc. 2022;3(2):101366.

